# *STREPTOCOCCUS PNEUMONIAE* SEROTYPE 22F INFECTION IN RESPIRATORY SYNCYTIAL VIRUS INFECTED NEONATAL LAMBS ENHANCES MORBIDITY

**DOI:** 10.1101/2020.06.09.142034

**Authors:** Sarhad Alnajjar, Panchan Sitthicharoenchai, Jack Gallup, Mark Ackermann, David Verhoeven

## Abstract

Respiratory syncytial virus (RSV) is the primary cause of viral bronchiolitis resulting in hospitalization and a frequent cause of secondary respiratory bacterial infection, especially by *Streptococcus pneumoniae* (*Spn*) in infants. While murine studies have demonstrated enhanced morbidity during a viral/bacterial co-infection, human meta-studies have conflicting results. Moreover, little knowledge about the pathogenesis of emerging *Spn* serotype 22F, and especially the co-pathologies between RSV and *Spn* is known. Here, colostrum-deprived neonate lambs were divided into four groups. Two of the groups were nebulized with RSV M37, and the other two groups mock nebulized. At day 3 post-infection, one RSV group (RSV/*Spn*) and one mock-nebulized group (*Spn only*) were inoculated with *Spn* intratracheally. At day 6 post-infection, bacterial/viral loads were assessed along with histopathology and correlated with clinical symptoms. Lambs dually infected with RSV/*Spn* had higher RSV titers, but lower *Spn*. Additionally, lung lesions were observed to be more intense in the RSV/*Spn* group characterized by increased interalveolar wall thickness accompanied by neutrophil and lymphocyte infiltration. Despite lower *Spn* in lungs, co-infected lambs had more significant morbidity and histopathology, which correlated with a different cytokine response. Thus, enhanced disease severity during dual infection may be due to lesion development and altered immune responses rather than bacterial counts.

## Introduction

Respiratory Syncytial Virus is one of the leading causes of severe lower respiratory infection in infants under the age of five, leading to 600,000 deaths worldwide [1]. RSV is a member of the pneumoviridae family that infects most infants by the age of two years [2]. Although a mild to moderate upper respiratory tract infection is the most common form of infection, severe lower respiratory tract infection can develop leading to bronchiolitis that frequently leads to hospitalization and sometimes death [3]. Lower respiratory tract infection can also increase the susceptibility to secondary bacterial infection(s) leading to severe and life-threatening pneumonia [4]. *Streptococcus pneumoniae* (*Spn*) is one of the most common bacterial infections that occurs concurrently with respiratory viruses such as influenza and RSV [5, 6], but unlike influenza, less is known about common etiologies during a dual RSV/*Spn* infection. In fact, while secondary bacterial pneumonia is well known for influenza, clinical data suggests RSV can also often cause bacterial pneumonia that is not well recognized. RSV is associated with invasive *Spn* such as pneumonia in the young or immunocompromised [7–11]. Other studies have demonstrated that RSV is the greatest cause of pneumonia in infants with co-infection with *Spn* very common [12–14]. However, mechanisms associated with secondary *Spn* pneumonia in RSV infected children are not well known.

*Spn* is a Gram-positive facultative anaerobic bacterial pathogen that causes invasive disease including sepsis, meningitis, and pneumonia. Similar to RSV, *Spn* causes severe illness and presents with a higher incidence in both children and the elderly worldwide [15]. Pneumococcal pneumonia is one of the leading causes of bacterial pneumonia in children worldwide, responsible for about 11% of all deaths in children under the age of five (700,000-1 million every year). Most of these deaths occur in developing countries [16]. *Spn* vaccines are effective in reducing the incidence of pneumonia caused by the serotypes contained in the vaccine [17]. However, the emergence of non-vaccine serotypes and persistence of antibiotic-resistant *Spn* such as serotype 19A highlights the importance of more investigation into *Spn* pathogenesis and therapy. Since *Spn* plays an essential role in secondary bacterial infections following viral pneumonia or viral-bacterial co-infection [15, 18], animal modeling for understanding viral-bacterial co-infections is crucial to investigating therapeutics that combat both. Moreover, most studies have concentrated on influenza and *Spn* co-infections but mainly in murine models with few mechanistic studies done in humans other than the calculation of frequencies of coinfections with these two pathogens [19–21]. Despite the importance of RSV/*Spn* co-infections, far fewer studies in this area as compared to influenza/*Spn* have been done. Furthermore, less is known about emergent serotype 22F pathogenesis [18, 22]. We have extensively used a neonatal lamb model to mimic RSV lower respiratory tract infection in infants as a preclinical model to evaluate the efficacy of new therapeutics [23] and to understand RSV pathogenesis [24–26]. Sheep are also permissive to *Spn* infection and have served as a model of *Spn* sepsis that appears to manifest clinical signs similar to human infection [27, 28]. Thus, our current study had a few objectives: (1) Can we model RSV/*Spn* pneumonia in a large infant animal species that can be improved in future studies; (2) can we successfully dually infect lambs and do we get enhanced disease; (3) can we use the model to gain insights into mechanisms that enhance morbidity over the single pathogen control groups; and (4) can use infant lambs to study *Spn* pathogenesis? We hypothesized, based largely on influenza dual infections, that RSV and *Spn* infected lambs would exhibit higher viral and/or bacterial burdens when dually infected. However, here we determined that pathogen burdens did not correlate with levels of viral lesions or bacterial burdens but rather with different immune responses between groups.

## Material and methods

### Experimental Design

#### Animals

A total of 20, 2-3 day-old, colostrum-deprived lambs, were randomly divided into four groups with 5 animals per group: RSV only, RSV-Spn co-infection, Spn only, and uninfected control. Animal use was approved by the Institutional Animal Care and Use Committee of Iowa State University. All experiments were performed following relevant guidelines and regulations as set by regulatory bodies. For viral inoculations, infectious focus forming units (IFFU), where only replication competent virus is detected by antibody in limiting dilution assays, were utilized. Two groups were exposed to nebulized RSV M37 (1.27×107 IFFU/mL), as done previously [29, 30], on day 0. One of the RSV infected groups was inoculated intratracheally with 2 ml normal saline as a mock Spn infection (RSV group) using syringe and needle, while the second RSV-infected group was inoculated intratracheally with 2 ml solution containing Spn serotype 22F (2×106 CFU/ml) 3 days post-RSV nebulization (RSV-Spn group). The other two groups were exposed to nebulized cell-conditioned mock media containing 20% sucrose at day 0 and inoculated intratracheally with either normal saline (control group) or solution containing Spn (2×106 CFU/ml) at day 3 post nebulization (Spn group). At day 6 post-RSV infection, all lambs were humanely euthanized with Fatal Plus. An autopsy was performed to evaluate the macroscopic lung lesions. After removal, each lung was examined by a pathologist similar to prior studies [30, 31]. If lesions were present, percentage involvement was estimated for each lung lobe. Percentages were converted to a scale using the following formula: 0%=0, 1-9%=1, 10-39%=2, 40-69%=3, 70-100%=4. Group averages were calculated for the gross lesion score. Lung samples were collected including sterile lung tissue for bacterial isolation, frozen lung sample for RT-qPCR, bronchioalveolar lavage fluid (BALF) from right caudal lung lobe for RSV IFFU assay and RT-qPCR, and lung pieces from different lobes were fixed in 10% neutral buffered formalin for histological assessment. Animals were observed daily and scored (1-5 on severity) by blinded animal caretakers concerning clinical symptoms including wheezing, lethargy, coughing, nasal/eye discharge while also taking a daily rectal temperature.

#### Infectious agents

Lambs were infected with RSV strain M37, purchased from Meridian BioSciences (Memphis, TN, USA). This strain is a wild type A RSV isolated from the respiratory secretions of an infant hospitalized for bronchiolitis [32, 33]. M37 was grown in HELA cells and stored at −80°C in media containing 20% sucrose [29]. 6 mL of 1.27 x 10^7^ IFFU/ mL in media containing 20% sucrose or cell-conditioned mock media (also containing 20% sucrose) was nebulized using PARI LC Sprint™ nebulizers to each lamb over 25-30 minutes resulting in the total inhalation of about 3 mL by each lamb [29]. *Spn* serotype 22F was grown overnight at 37°C in Todd Hewitt media containing 2% yeast extract, 50 μg/ml of gentamicin, and 10% bovine serum. Colony forming units (CFUs) were calculated by OD600 with confirmation by dilution plating on Tryptic Soy Agar (TSA) plates with 5% sheep blood containing gentamicin.

#### Lung RSV viral and *Spn* bacterial titers

BALF collected from the right caudal lobe at necropsy by flushing the caudal lobe with 5 mL of cold DMIM and collected back several times as done previously [29, 30]. Collected BALF was used to evaluate RSV IFFU (Plaque assay that counts the number of syncial cells formed due to viral infection detected by fluid fluorescent antibody technique). BALF was spun for 5 minutes at 3,000g to pellet large debris. Supernatants were spun through 0.45 am Costar SPIN-X filters (microcentrifuge 15,600g) for 5 minutes. The resulting BALF samples were applied to HELA cells grown to 70% confluence in 12-well culture plates (Fisher Scientific, Hanover Park, IL) at full strength, and three serial dilutions (1:10, 1:100, and 1:1000); all samples were tested in triplicate to determine the viral titer. Plates were stained with fluorescent antibody technique and as described previously [29, 30]. 100 μL of the right caudal lobe BALF was added to 1 mL TRIzol (Invitrogen) and kept in – 80 °C for the qRT-PCR assay to assess RSV mRNA. Sterile lung tissue samples were used to determine *Spn* titer. Lung tissue samples were placed in 500 μl of sterile PBS and were mechanically homogenized by a pestle. Lung homogenates were pelleted at 100xg, for 5 minutes. Supernatants were serially diluted and applied to 5% sheep blood TSA plates containing gentamycin.

#### Immunohistochemistry (IHC)

Formalin-fixed paraffin-embedded tissue sections were used for IHC, which was performed according to a previously published protocol in our laboratory [26, 29]. Briefly, after deparaffinization and rehydration, antigen retrieval was performed in 10mM TRIZMA base (pH 9.0), 1mM EDTA buffer, and 0.05% Tween 20 with boiling under pressure for up to 15 minutes. Polyclonal goat anti-RSV antibody (Millipore/Chemicon, Temecula, CA; Cat. No. AB1128) was used as the primary antibody after two blocking steps. The first blocking was with 3% bovine serum albumin in Tris-buffered saline+0.05% Tween 20 (TBS-T), and the second was 20% normal swine serum in TBS-T for 15 minutes each. The primary antibody was followed by the application of a biotinylated rabbit anti-goat secondary antibody (KP&L; Cat. No. 16-13-06). Signal development was accomplished using a 1:200 dilution of streptavidin-horseradish peroxidase (Invitrogen; Cat. No. 43-4323) for 30 minutes followed by incubation with Nova Red chromagen solution (Vector; Cat. No. SK-4800). A positive signal was quantified in both bronchioles and alveoli for each tissue section, and a score of 0-4 was assigned according to an integer-based scale of: 0=no positive alveoli/bronchioles, 1=1-10 positive alveoli/bronchioles, 2=11-39 positive alveoli/bronchioles, 3=40-99 positive alveoli/bronchioles, 4=>100 positive alveoli/bronchioles. IHC for *Spn* was performed using rabbit *anti-Streptococcus pneumoniae* polyclonal antibody (Thermo Fisher scientific cat. # PA-7259) followed by biotin-labeled goat anti-rabbit IgG antibody (Thermo Fisher Scientific Cat.#: 65-6140). Five random images were taken for each tissue section that was then analyzed by the quantitative Halo program. **Quantitative reverse transcription polymerase chain reaction (RT-qPCR):** BALF and lung tissue homogenates in Trizol were used to assess RSV mRNA expression by RT-qPCR. The assay was performed as published previously in our laboratory **[26, 30, 31]**. Briefly, RNA isolation from lung tissue and BALF was performed using the TRIzol method followed by standard DNase treatment. RT-qPCR was carried out using the One-Step Fast qRT-PCR Kit master mix (Quanta, BioScience, Gaithersburg, MD) in a StepOnePlus™ qPCR machine (Applied Biosystems, Carlsbad, CA) in conjunction with PREXCEL-Q assayoptimizing calculations. Primers and probe for RSV M37 nucleoprotein were designed based on RSV accession number M74568. Forward primer: 5’-GCTCTTAGCAAAGTCAAGTTGAACGA; reverse primer: 5’-TGCTCCGTTGGATGGTGTATT; hydrolysis probe: 5’-6FAM-ACACTCAACAAAGATCAACTTCTGTCATCCAGC-TAMRA.

Additionally, PBMCs were harvested at 6 days post-infection and added to RNAlater (Sigma) and stored at −80 degrees after an overnight incubation at 4 degrees C. RNA was then isolated by an RNA plus isolation kit (Qiagen, Gaithersburg, MD) per the manufacturer’s directions and then subjected to qRT-PCR using a single step reaction using Luna reagent (NEB, IPswhich MA). The primers and probes (5’-6FAM and Iowa Black Quencher) used were for IL-10, IFN γ, Actin, IL-1β, and IL-17a designed using published lamb cytokine sequences and PrimerDesign (UK) to find optimal pairs. For the detection of changes in gene expression (normalized on Actin), the RNA levels for each were compared with the levels in uninfected lambs (calibrators), and data are presented as the change in expression of each gene. The Δ*C_T_* value for the tissue sample from the calibrator was then subtracted from the Δ*C_T_* value of the corresponding lung tissue of infected mice (ΔΔ*C_T_*). The increase in cytokine mRNA levels in lung tissue samples of the infected animals compared to tissue samples of baseline (calibrator) animals was then calculated as follows: increase = 2^ΔΔ*CT*^.

#### Hematoxylin-eosin staining and histological scoring of lung sections

Hematoxylin-eosin stained sections were examined via a light microscope. An integer-based score of 0-4 was assigned for each parameter (bronchiolitis, syncytial cells, epithelial necrosis, epithelial hyperplasia, alveolar septal thickening, neutrophils in bronchial lumen, neutrophils in alveolar lumen, alveolar macrophages, peribronchial lymphocytic infiltration, perivascular lymphocytic infiltration, lymphocytes in alveolar septa, fibrosis), with 4 as the highest score. A final score was calculated by adding up all measured scores to form a 0-48 score, with 48 as the highest, which is called the accumulative histopathological lesion score.

#### Dual co-localization studies

Hela and Vero cells were infected with RSVA2 (MOI of 0.05) expressing mKate2 fluorescent reporter for 24 hours. Media was washed and replaced with DMEM without antibiotics and labeled *Spn* (serotypes 6c, 19A, and 22F) similar to (Verhoeven et al., 2014) was added for an additional 4 hours at 37 degrees before washing with PBS and fixing using 2% paraformaldehyde. A Zoe fluorescent microscope was used to randomly document both pathogens on the cells in at least 10 fields with all setting similar overlapping the red and green channels on the brightfield.

#### RSV infection of Sheep neutrophils

Sheep neutrophils were obtained by Ficoll gradient centrifugation with removal of PBMCs. Neutrophil/blood pellets were then lysed in ACK lysis for 5 minutes on ice followed by washing in PBS. Neutrophils were then resuspended in DMEM 10% and infected with RSVa 2001 at MOI of 1 for 4 hours. Neutrophils were then washed 3 times and held in RNAlater until qRT-PCR for RSV F transcripts could be performed.

#### Statistical analysis

Statistical analysis used the Wilcoxon signed-rank test for nonparametric parameters such as accumulative microscopic lesion scoring, followed by nonparametric comparisons for each pair also using the Wilcoxon method. One-way ANOVA was followed by all pairs comparison by the Tukey-Kramer HSD method for gross lesion scores and viral titer analyses by RT-qPCR and IFFU assays.

## Results

### Infected lambs replicated RSV and were premissive for *Spn* infection

RSV titers and *Spn* colony-forming units were measured in this study to evaluate the degree of infection by each pathogen and to investigate the possible effect(s) of co-infection in the combined RSV-*Spn* group on the replication of each infectious agent. As measured by IFFU, infectious RSV was detected in both RSV and RSV-*Spn* groups. Although not significantly different, RSV titer was about 2 fold higher in RSV-*Spn* group (Figure 1a). A similar trend was observed when assessing RSV RNA detected in BALF by RT-qPCR (7.28 and 7.31 viral genomes/ml) (Figure 1b). Furthermore, similar to the viable virus titer increase, RSV virions measured by RT-qPCR in the lung of the RSV-*Spn* group were 2 fold higher than the RSV-only group, but again not significantly different (Figure 1c). While significant differences between groups were not detected with respect to viral burdens, bacterial burdens did exhibit some differences. Specifically, *Spn* was isolated in the lung tissue of both the *Spn* only and the RSV-*Spn* groups with the bacterial titer 8.3-fold higher in the *Spn* only infected group (p<0.05) (Figure 1d).

**Figure 1.**
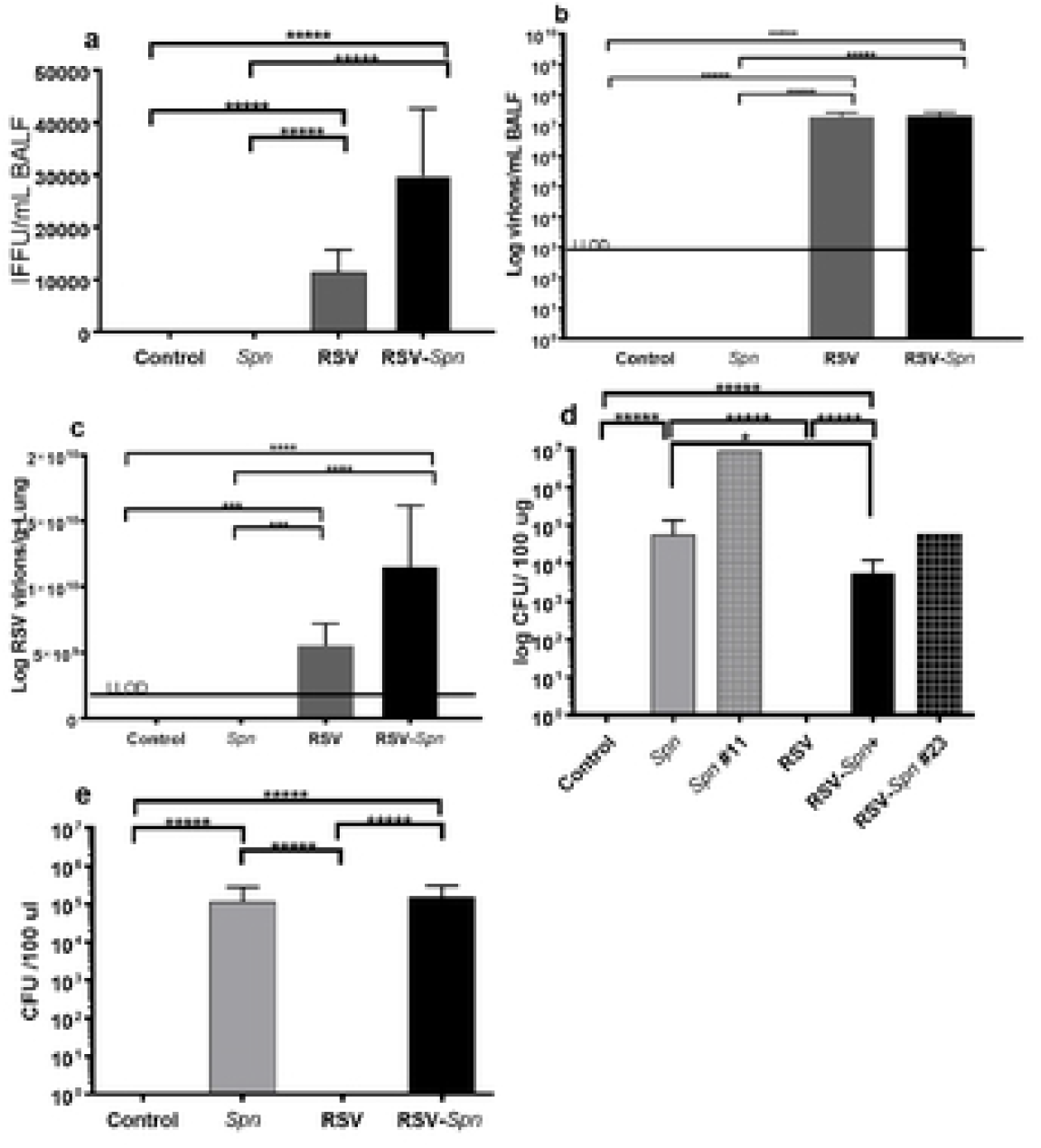
RSV and *Spn* titer in lung tissue and blood. (a) Number of infectious RSV particles as measured by IFFU assay, (b) RSV mRNA level in the BALF, (c) RSV mRNA level in lung tissue (d) *Spn* colony forming unit per 100 μg lung tissue, (e) *Spn* colony forming unit per 100 μl blood, all shown as average + SEM. Animals were either infected with mock media (control), RSV, *Spn*, or RSV followed by *Spn* (RSV-Spn). #11 and #23 died 48 and 36 hrs after *Spn* inoculation. *P<0.05, **P<0.01, ***P<0.005, **** P<0.001, ***** P<0.0001.

Interestingly, *Spn* titers in lambs that died before the end of the study were the highest of their groups. One lamb in the RSV-*Spn* group was found dead 36hr after bacterial inoculation with 59,302 CFU/μg in the lungs determined, while another lamb in the *Spn only* group was euthanized 48 hr after bacterial inoculations due to humane end-point being reached and had a titer of 9,302,325 CFU/μg (Figure 1d). Both of these animals had much higher bacterial counts then their group peers possibly indicating some loss of innate control over the bacteria. Unfortunately, *Spn* was detected in the blood in both *Spn* infected groups indicating bacteremia/sepsis (Figure 1e) development and possibly indicating a need for further model refinement (i.e. CFU given or route).

### Dually infected lambs showed elevated morbidity over *Spn* only

Daily temperatures were taken from each lamb during the study. While uninfected and RSV groups failed to spike a temperature at any point during the infection, *Spn* and RSV-*Spn* both exhibited an increase in body temperature after inoculation of the bacteria indicative of a mild fever as would be typical of *Spn* pneumonia (Figure 2A). However, the differences were not statistically significant, and both had similar temperatures 3 days post inoculation with the bacteria.

**Figure 2.**
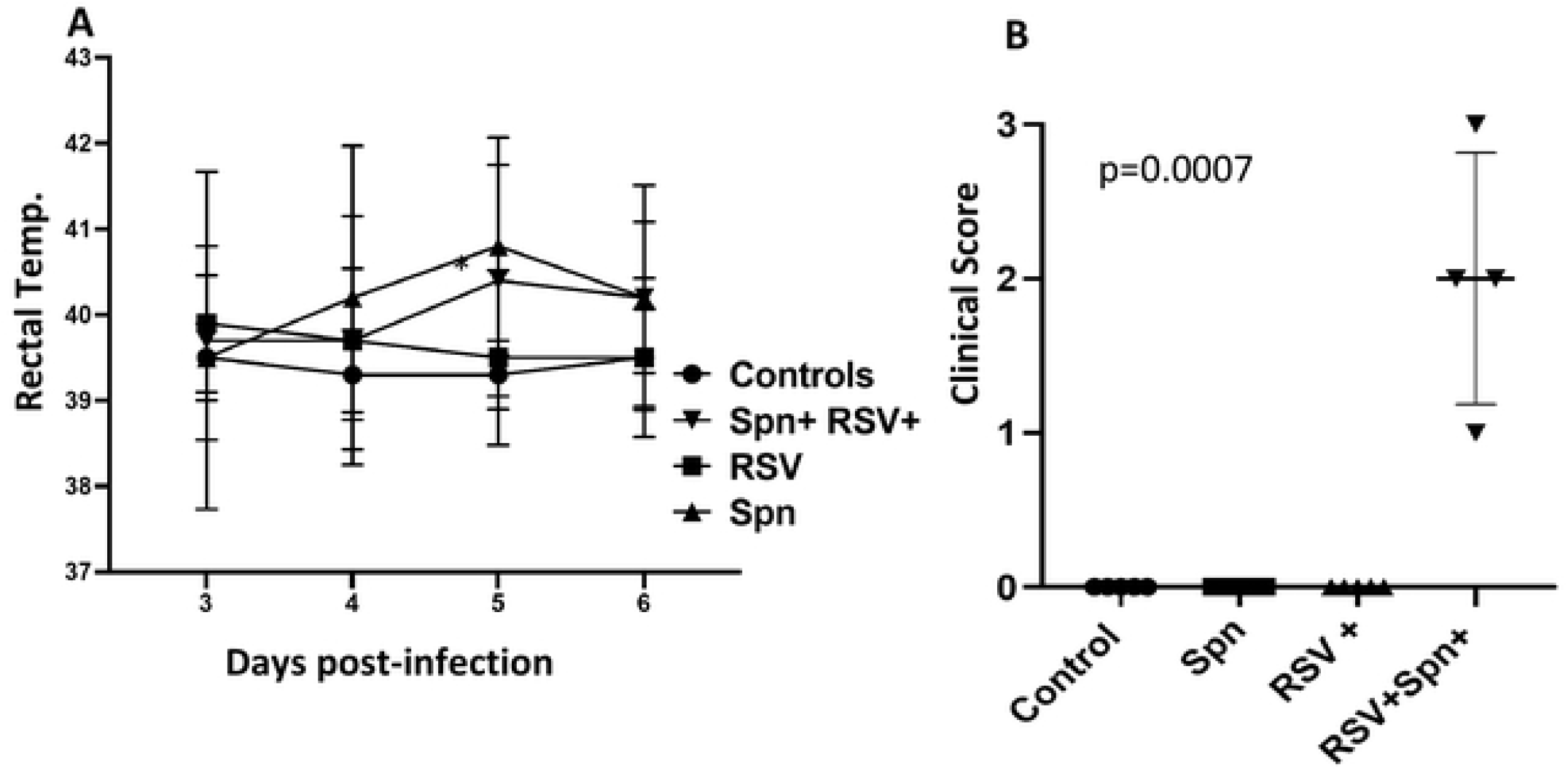
Morbidity levels after infection. (a) Rectal temperatures measured during infection, (b) level of morbidity observed (coughing, wheezing, lethargy, respiratory rate) rated from 1 (least severe) to 5 (most severe) as evaluated by blinded observation of the animals at 6 days post-infection.

Where the two groups did diverge was in the clinical symptom scores. By two days postinfection, only 3 of the 5 RSV-*Spn* lambs were scored by blinded animal care staff as visibly sick while only 1 of the 5 *Spn* only lambs was scored sick and that animal subsequently died from sepsis that day (data not shown). By three days post-infection, the *Spn* alone group still were not scored as showing symptoms while the RSV-*Spn* lambs all exhibited lethargy, coughing, or wheezing (Figure 2B).

### RSV and *Spn* induce a well-recognized macroscopic and microscopic lesion

Percent of lung tissue with gross lesions related to either infectious agent was determined at necropsy coupled with post-necropsy retrospective qualitative analyses. Both RSV and *Spn*-related lesions were found scattered across the lung surface in all lung lobes. Pinpoint dark red areas of lung consolidation characterized RSV lesions. These areas were evident in RSV and RSV-*Spn* groups. There were no differences in the percentage of the lung with RSV macroscopic lesions detected between RSV and RSV-*Spn* groups (Figure 3a &3c). *Spn* gross lesions are characterized by larger sizes of lung consolidation with bright red color - which was seen to a lesser extent when compared to RSV lesions (Figure 3b & 3d). There was a significant increase (p<0.001) in the percent of gross lesion in the RSV-*Spn* group since it has both RSV associated lesion and *Spn* associated lesion (Figure 3e).

**Figure 3.**
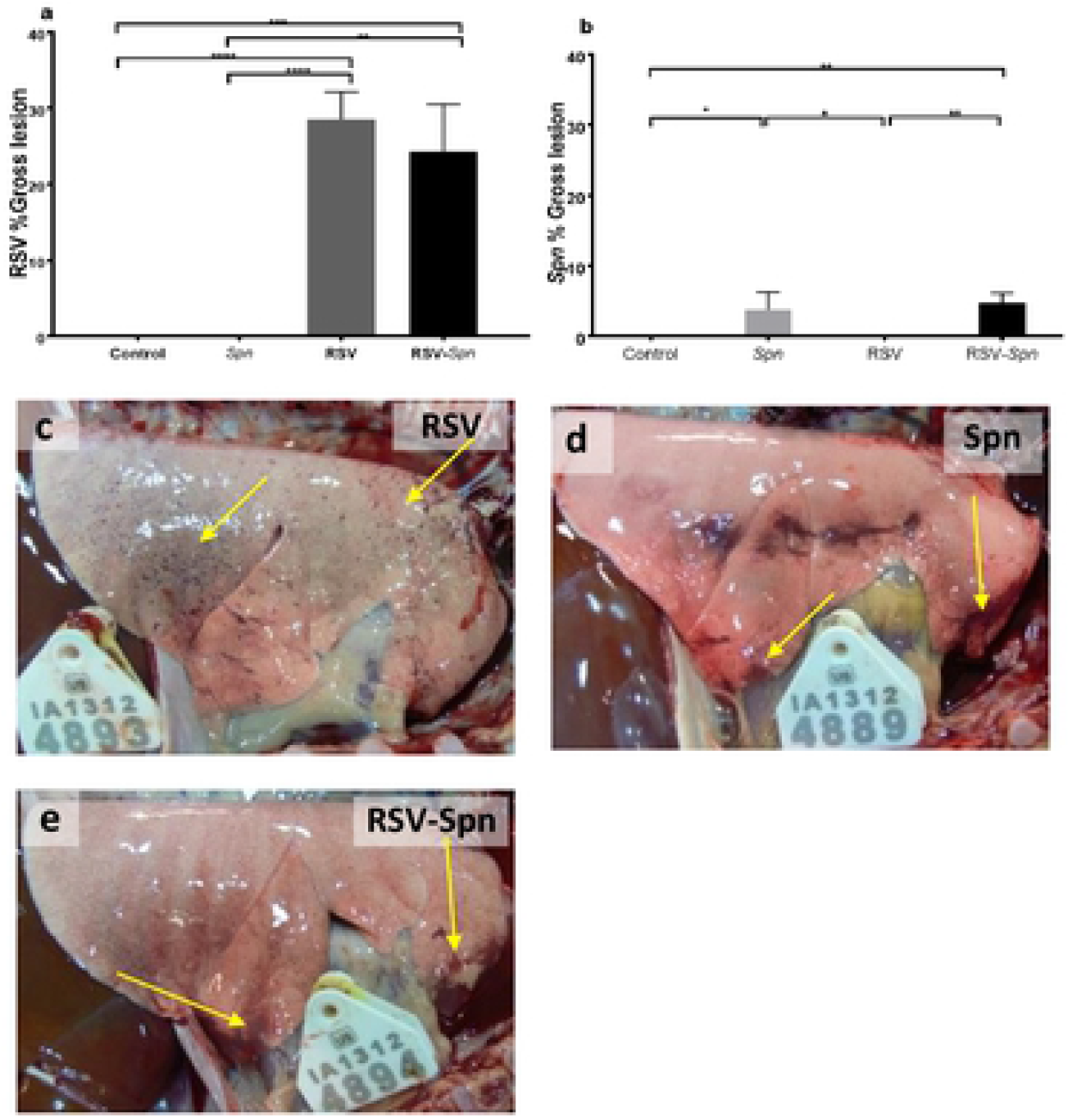
Percent of lung tissue associated with RSV and/or *Spn* infection. Percent of lung tissue associated with RSV lesions (a), *Spn* (b), with photographic representation of RSV-only (c), *Spn*-only (d), RSV-*Spn* (e). All show average and SEM. Lambs were either infected with mock media (control), RSV, *Spn*, or RSV followed by *Spn* (*RSV-Spn*). *P<0.05, **P<0.01, ***P<0.005, **** P<0.001, ***** P<0.0001.

Microscopic lesions observed within the lung tissue reflected the infectious agent used and contradicted our initial expectations (*i.e*., microscopic lesions caused by RSV infection were multifocal areas of interstitial pneumonia, and bronchiolitis scattered randomly and homogeneously throughout the lung tissue). However, *Spn* induced diffuse homogenous and subtle pathological changes in the lung tissue. Infection with either *Spn*, RSV, or both, markedly increase microscopic lesions (accumulative microscopic lesion score) associated with the disease in comparison to the control group (p<0.05) (Figure 4a-f). Additionally, the combined RSV-*Spn* infection significantly increased the severity of microscopic lesions in comparison to the *Spn* only group (p<0.05). Lesions varied among lambs, and RSV lesions consisted of thickening of the interalveolar wall with inflammatory cellular infiltrates in the airway adventitia and lamina propria (lymphocytes and plasma cells), the alveolar lumen (alveolar macrophages and neutrophils), and bronchiolar lumen (neutrophils). With RSV, overall, there was a varying degree of epithelial necrosis and syncytial cell formation. On the other hand, *Spn* lesions consisted of moderate interalveolar wall thickening with inflammatory cellular infiltrate, mainly in the alveolar septae. Most of the microscopic lesions seen with RSV overlapped with *Spn*-induced injury. However, congestion of the interalveolar wall capillaries and hemorrhage was seen only in *Spn*-inoculated lambs.

**Figure 4.**
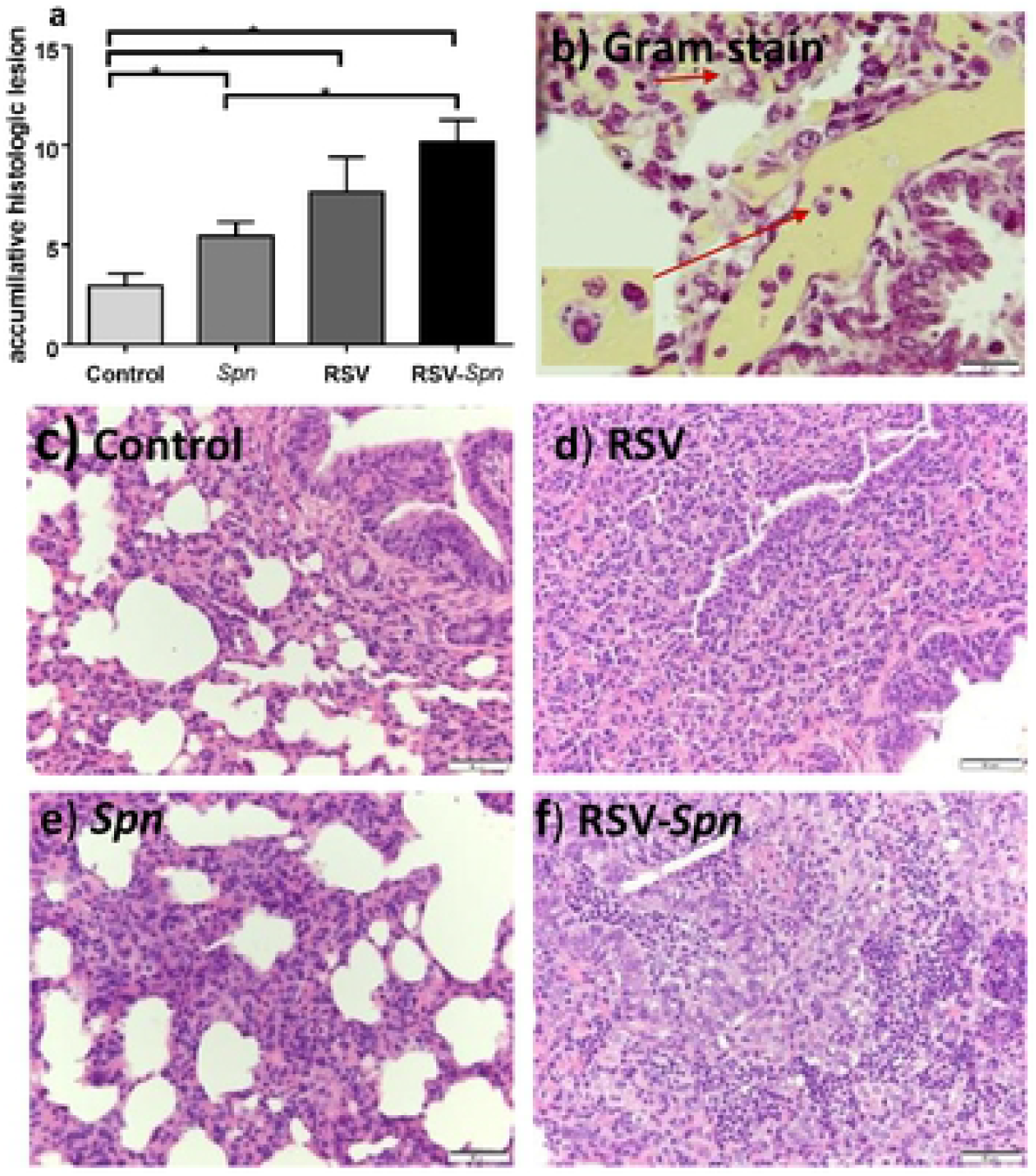
Histologic lesions associated with RSV, *Spn*, and RSV-*Spn* combined infection. (a) Accumulative histologic lesion associated with RSV and *Spn* infection shown as average + SEM (b-f) show a representative photograph of lung tissue sections stained with Gram stain (b), H&E stained tissue section of control (c), RSV only (d), *Spn* only (e), combined *RSV-Spn* (f). Lambs were either infected with mock media (control), RSV, *Spn*, or RSV followed by *Spn* (RSV-*Spn*). * P<0.05, **P<0.01, ***P<0.005, **** P<0.001, ***** P<0.0001.

Immunohistochemistry was used to identify and localize RSV and *Spn* in tissue sections. RSV was present multifocally throughout the sections with bronchial and peribronchial distribution (Figure 5c). Therefore, RSV expression was evaluated in bronchioles and alveoli separately. There were no significant differences between the RSV only and RSV-*Spn* groups in the degree of RSV expression in lung tissue sections (Figure 5a). *Spn* was random and homogenously scattered throughout the lung sections with more intense signals in interalveolar walls and blood capillaries (Figure 5d). Although not significant, there was a 1.5 fold increase in *Spn* expression in the *Spn* only group when compared with the RSV-*Spn* group (Figure 5b).

**Figure 5.**
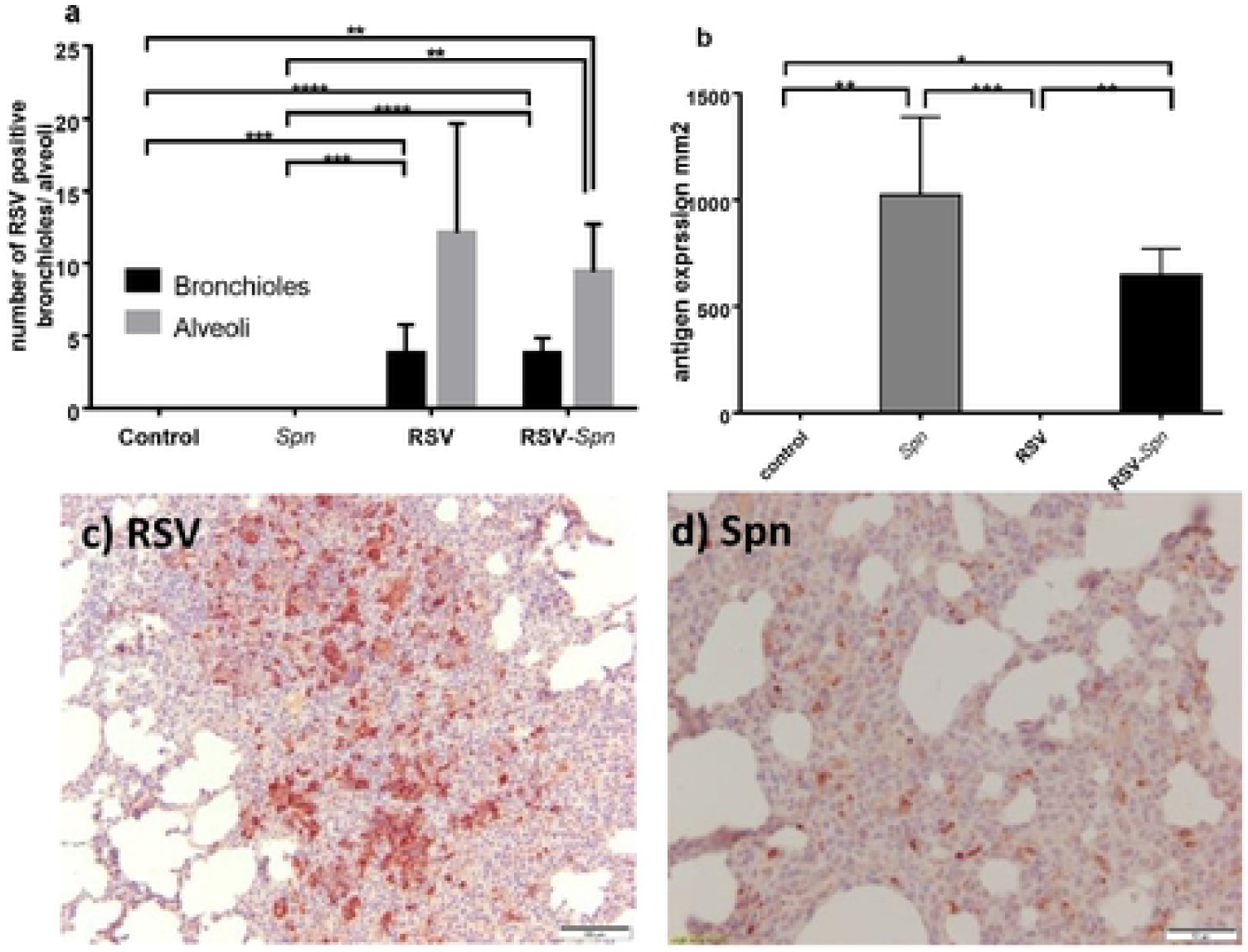
Immunohistochemistry staining of RSV and *Spn* in FFPE lung tissue sections. The number of bronchioles and alveoli express the RSV positive signal (a), surface area (mm2) occupied by *Spn* IHC positive staining (b), all shown as average + SEM, 5 fields examined. (c) and (d) show a photo representation of RSV (c) and *Spn* (d) IHC positive staining. Animals were either infected with mock media (control), RSV, *Spn*, or RSV followed by *Spn* (RSV-*Spn*). * P<0.05, **P<0.01, ***P<0.005, **** P<0.001, ***** P<0.0001.

### Divergent cytokine responses occurred between the groups

We next examined cytokine responses in PBMC from lambs at necropsy and found some significant differences in patterns by qRT-PCR between groups (Figure 6). Using the uninfected controls as the baseline, we found that PBMCs from the RSV only group were positive for IFNγ, IL-1β while *Spn* only group were positive for IL-10, IFNγ, and IL-1β. In contrast to both of these groups, RSV-*Spn* lambs were positive only for IL-1β. No IL-17a was detected in any of the lambs, and IL-4 was detected only in the uninfected controls.

**Figure 6.**
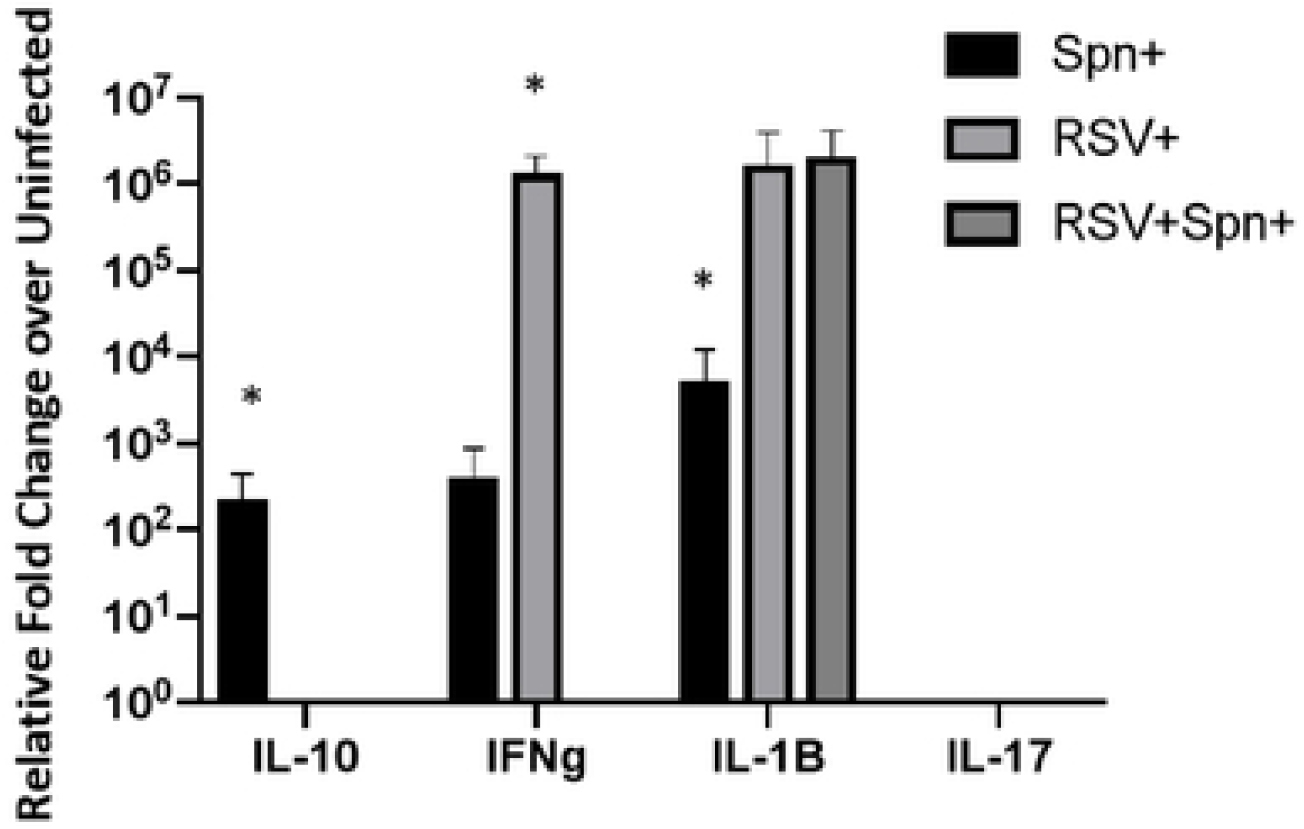
Cytokines in peripheral blood lymphocytes. qRTPCR was performed on isolated PBMCs at 6 days post-infection and shown as fold change over uninfected controls. **Figure 7. Dual infections of Hela and Vero cells.** RSV infection of Hela and Veros proceeded incubation with FITC stained RSV 19A and 22F. Fluorescent microscopy was used to examine for co-localization of both pathogens.

### Dual infections in Hela and Vero cells failed to demonstrate enhanced *Spn* attachment to infected cells

Since the literature suggests that Hela cells allow for intact G protein on virions while Vero cells cause a cleavage and the G protein is thought to directly bind to *Spn [34–36]*, we thought to explore these mechanisms since we found many of our *Spn* lesions may not have overlapped with RSV lesions. However, as shown in figure 7, we failed to observe enhanced RSV and *Spn* dual binding to either Hela or Vero cells suggestive that these two pathogens may not necessarily be interacting as we observed in lung lesions of the lambs. In fact, all three serotypes tested, while many co-localized with RSV infected cells, appeared to have an equally likelihood of attaching to non-RSV infected cells. Hela and Vero cells made no difference in any of these results.

**Figure 7.**
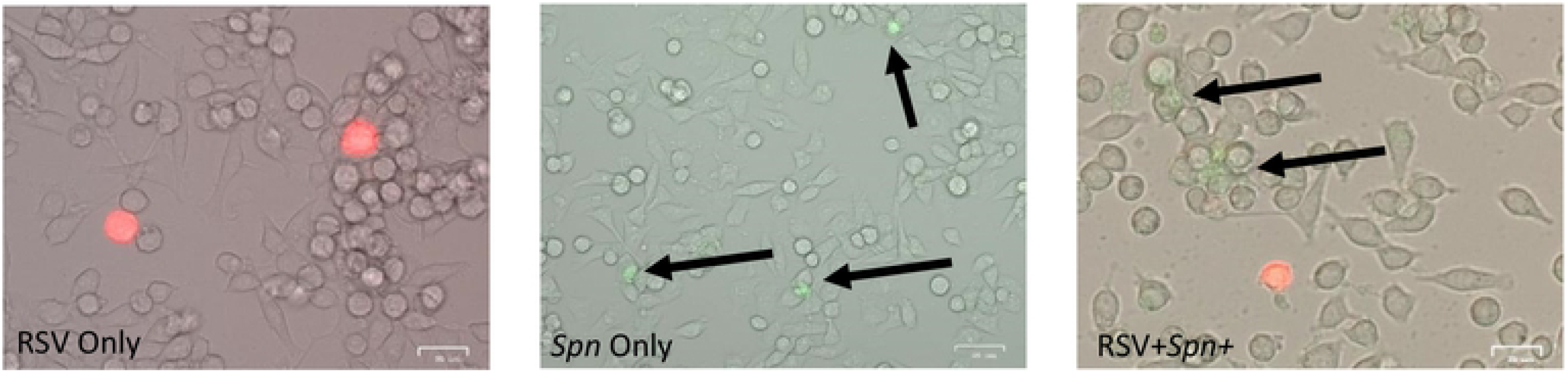
No strong association between infected cells and *Spn* attachment.

### RSV infection of lamb phagocytes are permissive

We next sought to determine whether infection of neutrophils could be occurring in our model and perhaps increasing their pathologic response in the lungs. In prior studies, we found that human infant neutrophils could be infected and this disrupted their in vitro *Spn* phagocytic activities. Furthermore, infants with severe RSV infections have been observed to have infected blood white blood cells. Thus, we obtained sheep neutrophils and infected them with 1 MOI of RSVa 2001 virus and allowed the infection to occur for 4 hours prior to extensive washing and examination for RSV F transcripts by qRT-PCR. Similar to human neutrophils, we found that sheep neutrophils could be infected with the virus (Figure 8A). In vivo staining for RSV also demonstrated many monocytes/macrophages infected by RSV (Figure 8B) which could also change their activities toward *Spn* in the lungs and also worthy of follow-up studies.

**Figure 8.**
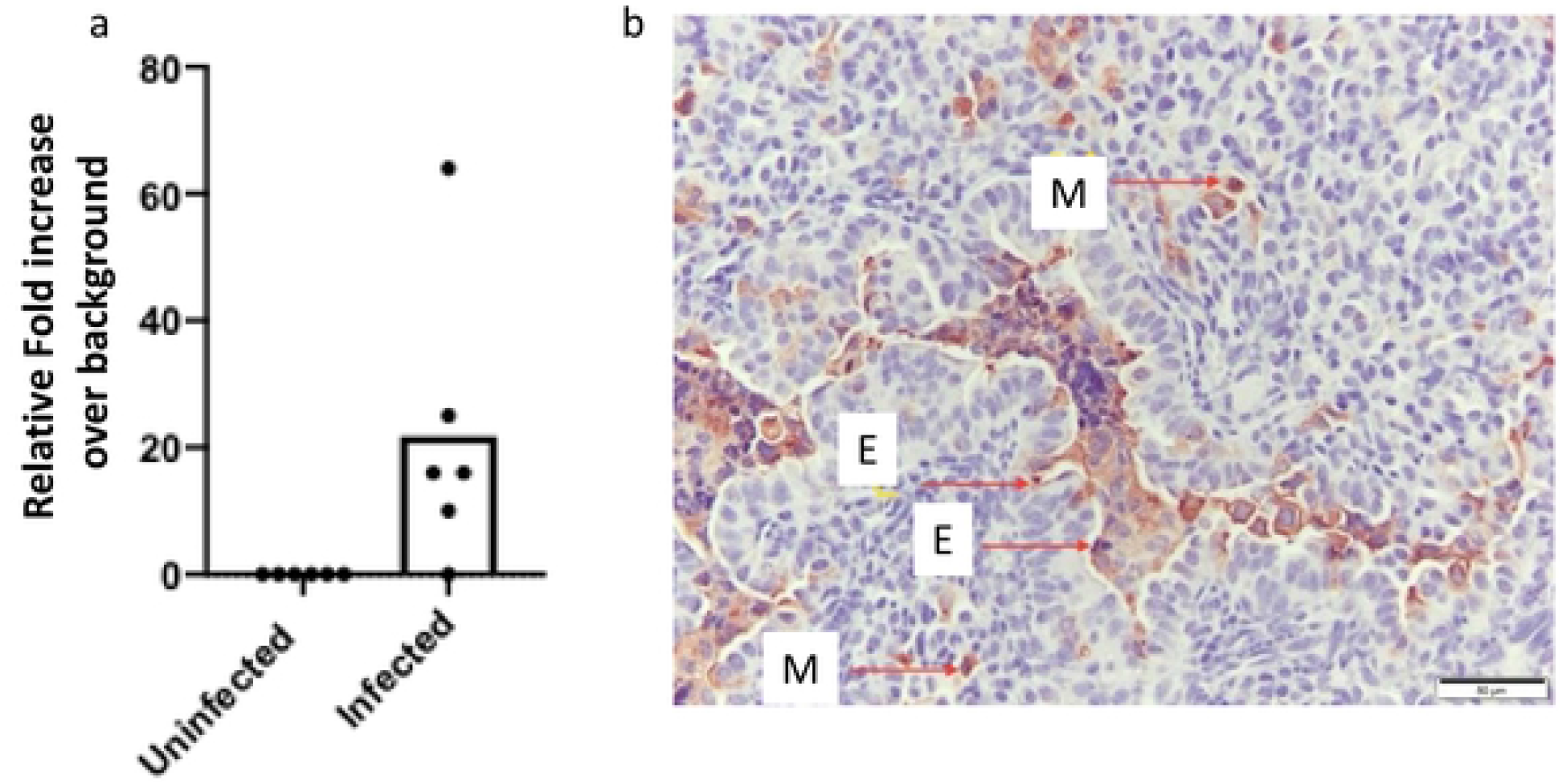
RSV infects phagocytic cells. (a) RSV infection occurs in sheep neutrophils as determined by qRT-PCR after infection of peripheral blood neutrophils in vitro, (b) RSV immunohistochemistry shows many infected monocytes/macrophages in the lungs of infected lambs. E=epithelial cell, M=monocyte/macrophage

## Discussion

There is a critical need for an animal model to study bacterial pneumonia secondary to an initial viral infection in the lung to study the mechanisms of viral-bacterial co-infection and to evaluate therapeutic interventions. There are significant advantages of using lambs to model RSV infection as a correlate for human infants - including the ability to use human viral strains without adaptation and the similarity of the pathological sequelae [24, 25]. Serotype 22F is thought to have appeared after the introduction of the *S. pneumoniae* vaccine PCV7 ([37]). While this serotype was not widespread in those subsequent years ([37]), 22F is now the second most common serotype causing invasive disease in children less than seven years old and the primary cause in the elderly [38]. Molecular analysis of this serotype also indicates six different lineages and 18% of clinical isolates demonstrating erythromycin resistance ([18]). Thus, this emerging serotype is a component of the PPSV23 vaccine and an important pathogen to observe for in children vaccinated with PCV13.

In this study, we demonstrate that *Spn* readily infects the lungs of lambs and establishes active bacterial pneumonia. A previous study revealed that the peak of RSV titer and infection in lambs is around day 3 post-viral nebulization, and we used this time-frame to model early human coinfection [30]. The results of this study demonstrate consistency in the infection rate of both RSV and *Spn*, as well as an excellent relation to the lesion development induced by either of the infectious agents. Although we used 2 × 10^6^ CFU of *Spn* for infection, murine studies typically use 5×10^5^ to 10^7^ CFU to get productive infections. Moreover, the lung volume of lambs is significantly larger than mice, which suggests that our inoculating dose may be more dispersed throughout the lungs than murine studies. We believe that we may also be able to reduce the infection dosage to a lower CFU or potentially use a colonization model to examine co-infection and pneumonia development.

Prior studies in mice and cotton rats with influenza or RSV/*Spn* co-infections demonstrated higher viral loads in dually infected animals [39, 40], although our observed viral (RSV) was not different in this study. Influenza co-infection studies also predict higher *Spn* burdens in the lungs due to damaged epithelial cells serving as anchor points for the opportunistic bacteria. In other studies, RSV with *Spn* in mice or cell culture predicts that the RSV G protein on the infected epithelial surface could also serve as an anchor point for *Spn* in the lungs [34]. However, we did not observe this in infection of either Hela or Vero cells. In contrast to these murine models, we found lower bacterial loads in the co-infection group over the *Spn* only group. These findings suggest that the immune response might control *Spn* in the lungs of lambs better than mice. Importantly, in human clinical studies of co-infection, an increase in nasal colonization numbers of *Spn* upon viral infection has been demonstrated, but this does not translate into higher invasive lung disease [41]. These suggest that higher bacterial burdens could be a murine artifact rather than a mechanism enhancing disease. In human studies of high *Spn* colonization, RSV disease appeared less severe [42], suggesting that further using the lamb model to explore mechanistic differences between *Spn* colonization and pneumonia during RSV. Of further interest, murine studies using IFNγ or IFNγ receptor knockouts and *Spn* infection have shown reduced lung CFUs over wild-type controls with no change in the level of morbidity [43]. Thus, the lower *Spn* counts that we observed in our RSV-*Spn* group could derive from the limited IFN γ response observed in these lambs. Six days post-infection is early for the recruitment of T-cells into the lungs, with three days post-*Spn* also much too early for antibacterial T-cells to infiltrate the lungs. However, peripheral blood could have early trafficking PBMCs migrating between lymph nodes toward the lungs. We are not yet sure why we observed high levels of IFN γ in the RSV group but not the RSV-*Spn* group, but it is feasible that the presence of the bacteria after the virus changed the character of the antiviral T cell response.

The only deaths that occurred in the present study were in the *Spn*-infected groups, and both lambs (lamb 11 in the *Spn* only group, and lamb 23 in the RSV-*Spn* group) had high lung *Spn* colony-forming units/gram tissue. These could represent a failure to control bacterial division and subsequent septicemia.

Lesion severity was consistent with the RSV titer and *Spn* burden as is shown by the significant increase in the percent of lung tissue involved by gross lesions, and the increase in the evaluated histological parameters. RSV gross lesions were multifocal lesions scattered randomly in all lung lobes - which is the typical lesion distribution induced by RSV nebulization [26, 30]. However, presentation of *Spn* gross lesions contradicted what was expected by the apparent development of lesions in all lobes - including the caudal lung lobe, which is not typical for bacterial pneumonia in lambs. However, the diffuse bacterial lesions and the presence of *Spn* lesions in the caudal lobe may be due to the inoculation technique used for *Spn* infection. For *Spn* infection, lambs were held vertically by one person and injected intratracheally by the second person leading to a fall of inoculum through the bronchial tree into the caudal lobe, which in this case, was favorable since it gives a bronchopneumonic distribution similar to that found in humans. It is also possible that *Spn* spreads across lung lobes after inoculation either by airflow or vascular flow. RSV-induced microscopic lesions were more prominent in comparison to *Spn-* induced lesions and subsequently led to significant differences between the RSV-*Spn* and *Spn* only groups’ accumulative histologic lesion scores. RSV was more prominent in the bronchioles, while *Spn* was diffuse throughout the lung sections.

Although we are still evaluating mechanisms, we believe that the higher morbidity observed in the RSV-*Spn* group may derive from an enhanced neutrophil response found in the lungs. Evidence for this was found in the histopathology and the lower *Spn* burdens in these animals. Likely, RSV infection served as a first activating response to neutrophils that could have then better controlled the secondary bacterial infection. It is also possible that alveolar macrophages were activated by RSV that, in turn, secreted inflammatory mediators that enhance neutrophil activation. Enhanced neutrophil/leukocyte activation contrasts with studies in influenza co-infections in mice – which suggests innate immune exhaustion [44]. While the time of inoculation could be a reason for the observed differences, another could be the small difference between influenza and RSV pathogenesis. In either case, the results suggest further avenues of study using this model. Of interest, morbidity is highest in infants infected with RSV that exhibit significant wheezing [45], and here we observed high wheezing in the presence of dual infection which further suggests that some of the increased morbidity could be from altered immune responses over the viral only group. The observed higher RSV infection rate in the co-infection could also derive from the higher number of neutrophils in the lungs in this group. There is evidence that RSV can infect neutrophils in humans [45], including our unpublished data. Thus, if dual infection with *Spn* leads to enhanced neutrophil recruitment to the lungs over RSV alone, those cells could become infected and contribute to the higher viral titer we observed in the dual infection group. The effects of RSV infection on neutrophilic antibacterial responses would be an interesting further study.

In this study, we have developed an animal model of co-infection for RSV and *Spn*. Of course, limitations in the study include low sample numbers in each group that may have limited our ability to achieve some statistical differences in RSV titers. However, prior studies by the authors were adequately powered at these sample numbers. We have determined enhanced disease with co-infection of both pathogens that mirrors studies of human and murine influenza infections, but this may all be due to a complex enhanced inflammatory/immune response to co-infection rather than direct damage by either pathogen alone. Additional studies will allow refinement of this model and will include variations in inoculum volume/concentration, the time between infections, and kinetic analyses.

## Acknowledgments

The work was sponsored by an investigator-initiated grant awarded to DV by Merck.

## Author contributions

All authors were involved in animal infections, necropsies, data analysis, and manuscript writing. SA also performed immunohistochemistry, pathologic analysis of histology, prepared viral stocks, and had a primary role in the manuscript writing. PS helped prepared tissues for histology and helped with the analysis of histology. JG performed qRT-PCR for viral loads. MA performed the primary pathologic and immunologic analysis of lung histology. DV performed bacterial preparation, lung CFU counts, cytokine analysis, and experimental design.

## Data Availability

The datasets generated and analyzed during the current study are available from the corresponding author on reasonable request.

